# Upregulating action semantics with neuromodulation and gesture observation to facilitate verb retrieval in aphasia

**DOI:** 10.64898/2026.04.10.716321

**Authors:** Haley C. Dresang, Laurel J. Buxbaum, Roy H. Hamilton

## Abstract

Although many individuals with chronic aphasia respond to language therapy, there remains a need for adjunctive interventions that can enhance treatment response. Approaches targeting multiple modalities, such as gesture cueing, and neuromodulation techniques, such as transcranial magnetic stimulation, have shown promise for supporting language recovery. The present pilot study investigated whether enhancing activation of the action semantic network could facilitate verb production in individuals with chronic aphasia. Participants were recruited as a convenience sample and completed a within-subject design in which all individuals received each condition. Two non-linguistic methods of activating the action semantic network were evaluated: (1) pantomimed gesture cues to prime action concepts and (2) intermittent theta-burst stimulation to the left posterior middle temporal gyrus (pMTG), an intact action-semantic network node in our participants. We examined individual and combined effects of gesture priming and stimulation to test whether a combined approach would yield additive or interactive benefits. Using a Bayesian generalized linear mixed-effects model, we observed a moderate interaction between gesture priming and stimulation site. Contrary to predictions, combining gesture priming with pMTG stimulation did not produce additional benefits over either intervention alone. Instead, pMTG stimulation attenuated the priming advantage observed under vertex stimulation, and gesture priming attenuated the advantage observed with pMTG stimulation alone. Posterior estimates provided substantial preliminary evidence for this interaction in our pilot sample size. These findings suggest that combined activation of the action semantic network through gesture and neuromodulation approaches may not benefit verb retrieval above and beyond each approach alone.

## INTRODUCTION

Aphasia is a disorder of language that is experienced by approximately one in three stroke survivors (Berthier, 2005). One robust predictor of everyday communicative function and quality of life is the degree of verb production impairment (Rofes et al., 2015), a challenge experienced by 70% of people with aphasia (Mätzig et al., 2009). Developing treatments for these impairments has implications for those living with aphasia after stroke as well as individuals experiencing verb impairments in the context of other neurological disorders such as Parkinson’s disease and Amyotrophic Lateral Sclerosis (Antonucci & Reilly, 2008; Bak, 2013; Bocanegra et al., 2017; Cardona et al., 2013; Cousins et al., 2018; York et al., 2014).

Several speech and language therapies that target verb-production impairments do so by strengthening lexical-semantic networks associated with a given verb. For example, the verb *chopping* can co-activate representations such as *chef* (agent), *carrot* (patient), *knife* (instrument), and *kitchen* (location; McRae et al., 2001, 2005). Training retrieval of semantically related words is posited to activate and strengthen the action semantic network, promoting generalized improvement in verb production, noun-verb combinations in sentence production, and widespread retrieval of untreated words (see Edmonds, 2016 for review).

Even with such verb-centered approaches, however, meaningful gains in chronic aphasia recovery often requires high-intensity speech and language therapy (e.g. Bhogal et al., 2003). Yet such intensity is rarely feasible in real-world clinical settings (Code & Petheram, 2011). Moreover, therapy outcomes vary due to intrinsic factors (e.g., lesion characteristics, cognitive profile) and extrinsic factors (e.g., time post-onset, therapy dose; Doogan et al., 2018). Although many individuals with chronic aphasia respond to standard language therapies, there remains a strong need for novel interventions that may improve treatment response and benefit additional individuals. As a result, there is growing interest in adjunctive approaches that employ multiple modalities, such as gesture cueing, and non-invasive brain stimulation, such as transcranial magnetic stimulation (TMS). The present study investigated the effects of gesture cueing and TMS to examine whether enhancing activation of the action semantic network can facilitate verb production in individuals with chronic aphasia.

Our central hypothesis was that pre-activating the action semantic network would facilitate accurate verb production. Action semantics is characterized theoretically as a broad conceptual system that encodes the meaning of actions (*chopping*), associated objects (*knife, carrot*), and events (*cooking*), and is accessible through multiple modalities, such as language, gesture, perception, and motor simulation (Lebkuecher et al., 2026; Reilly et al., 2025; van Elk et al., 2014). This semantic system is subserved by a distributed neural network, including the left posterior middle temporal gyrus (pMTG), inferior frontal gyrus (IFG), and posterior superior temporal sulcus (pSTS) (Holle et al., 2008, 2010; Vigliocco et al., 2020; Willems et al., 2007, 2009). These nodes and their connections are engaged during both verb retrieval (Dresang et al., 2021; Olson et al., 2025) and meaningful gesture processing (Dresang et al., 2023; Tarhan et al., 2015), suggesting a shared repository of action semantics. For instance, co-speech gestures and pantomimes activate left IFG and pMTG (Willems et al., 2009), and gesture-enhanced language elicits greater BOLD response in left pMTG and IFG than redundant gestures or language alone (Dick et al., 2014). Evidence from neurodegenerative disorders further implicates pMTG, pSTS, and IFG in verb but not noun production (Lukic et al., 2021), indicating that verbs and gestures access at least partially shared action semantic networks. Our study specifically examined whether gesture cueing or non-invasive brain stimulation could target this network to enhance verb retrieval in aphasia.

### Gesture cueing to activate action semantic networks

Gesture cueing involves using meaningful hand or body movements to support language comprehension and word retrieval. Observing gestures engages the action semantic system, potentially enhancing access to words. There is substantial evidence that verb production in individuals with aphasia can improve when clinicians perform congruent gestures during or prior to verb naming (Hanlon et al., 1990; Rodriguez et al., 2006; Rose, 2013). As a result, gesture cues have been incorporated into single-subject aphasia treatment studies (Boo & Rose, 2011; Marangolo et al., 2010; Raymer et al., 2012; Rodriguez et al., 2006; Rose & Sussmilch, 2008). However, many of these studies required, and did not control for, multiple processes that can be impaired following stroke, such as auditory comprehension and gesture production. These requirements limit generalizability.

Murteira and Nickels (2020) addressed several of these limitations by examining whether gesture observation alone could facilitate verb retrieval in aphasia. In a priming experiment, participants named action pictures that were preceded by congruent, unrelated, or no gesture videos. They found verb production was faster and more accurate following congruent gesture observation, suggesting semantic priming effects (see also Vigliocco et al., 2020). This pilot study adapted Murteira and Nickels’ (2020) paradigm to test whether adding neurostimulation to a gesture cueing protocol can further augment verb production by boosting activation within the action semantic network.

### TMS to activate action semantic networks

Non-invasive brain stimulation can modulate cortical excitability and promote neuroplasticity in perilesional and residual language networks (e.g., Hamilton et al., 2011; Turkeltaub et al., 2012). One form of repetitive TMS, intermittent theta burst stimulation (iTBS), has emerged as a promising modality because of its ability to induce excitatory effects in perilesional language networks (Chou et al., 2022; Szaflarski et al., 2018). When applied to left fronto-temporal regions, iTBS has been associated with improvements in verb naming, lexical retrieval, and other language functions in post-stroke aphasia (Allendorfer et al., 2021; Chou et al., 2022). Most TMS aphasia studies have employed inhibitory protocols and have targeted the right inferior frontal gyrus (rIFG), with the goal of reducing potentially maladaptive hyperactivity (Ren et al., 2014; Thiel et al., 2006; Wang et al., 2024). Few studies have used excitatory iTBS to upregulate residual left-hemisphere language nodes (Allendorfer et al., 2021; Chou et al., 2022).

The left pMTG is a strong candidate stimulation site because it plays a key role in lexical-semantic processing, verb retrieval, and action representation (Andric & Small, 2012; Hoffman et al., 2012; Lesourd et al., 2021; Lingnau & Downing, 2015; Peelen et al., 2012; Straube et al., 2012; Tarhan et al., 2015; Whitney et al., 2011). It has been proposed as a hub for the storage and integration of lexical and semantic information (Straube et al., 2012), playing a causal and supramodal role in integrating gesture observation with speech production (Zhao et al., 2018). Lesion studies further implicate left pMTG in semantic deficits across verbal and nonverbal tasks (Andric & Small, 2012; Straube et al., 2012), and damage to left pMTG may reduce the benefit of semantics on gesture performance (Dresang et al., 2023).

While iTBS shows promise for aphasia broadly, its effects on verb production and gesture cueing remain untested. Prior neurostimulation studies of verb retrieval have typically used inhibitory protocols to create transient virtual lesions in neurotypical populations (Cappa et al., 2002; Cappelletti et al., 2007) or have employed transcranial direct current stimulation (tDCS) to enhance naming by targeting motor cortex (Branscheidt et al., 2018; Meinzer et al., 2016) or IFG (Fiori et al., 2013; Marangolo et al., 2013). To our knowledge, no studies have combined gesture cueing with iTBS to enhance verb retrieval in aphasia. Since both gesture cueing and iTBS have independently improved language performance, we aimed to investigate whether focal excitatory stimulation of the left pMTG could augment the effects of gesture cueing on verb production.

### Current study design & hypothesis

We evaluated two non-linguistic methods of activating this network in individuals with chronic aphasia: (1) pantomimed gesture cues to prime action concepts, and (2) iTBS to left pMTG, an intact action-semantic node in our participants. We examined the individual and combined effects of gesture priming and pMTG stimulation, testing whether a combined approach yields additive or interactive benefits. As a foundational feasibility study, this work also aimed to estimate effect sizes and characterize variability in response to gesture priming and stimulation to guide future research. Given the high inter-individual variability in aphasia treatment response, we also report descriptive behavioral and neural measures, including baseline cognitive-linguistic performance and structural neuroimaging indices.

## METHODS

### Study Design

Participants underwent a total of four research sessions. In the first two sessions, we assessed baseline performance on a battery of tasks, including the experimental task (described below) without priming cues. These baseline sessions were conducted remotely due to the COVID-19 pandemic. In the following two sessions, subjects completed the experimental task with counterbalanced priming during cross-over sessions of excitatory iTBS to left pMTG and the vertex control site in pseudorandomized order. The behavioral experiment procedures and materials, adapted from Murteira & Nickels (2000), required participants to name action pictures using a complete sentence with a single verb phrase following the presentation of congruent or neutral gesture primes. Baseline assessment sessions were scheduled at least one day apart, and treatment sessions were scheduled at least one week apart to allow a washout period and help reduce practice effects. See Figure 1 for a diagram of the research session timeline.

**Figure 1.**
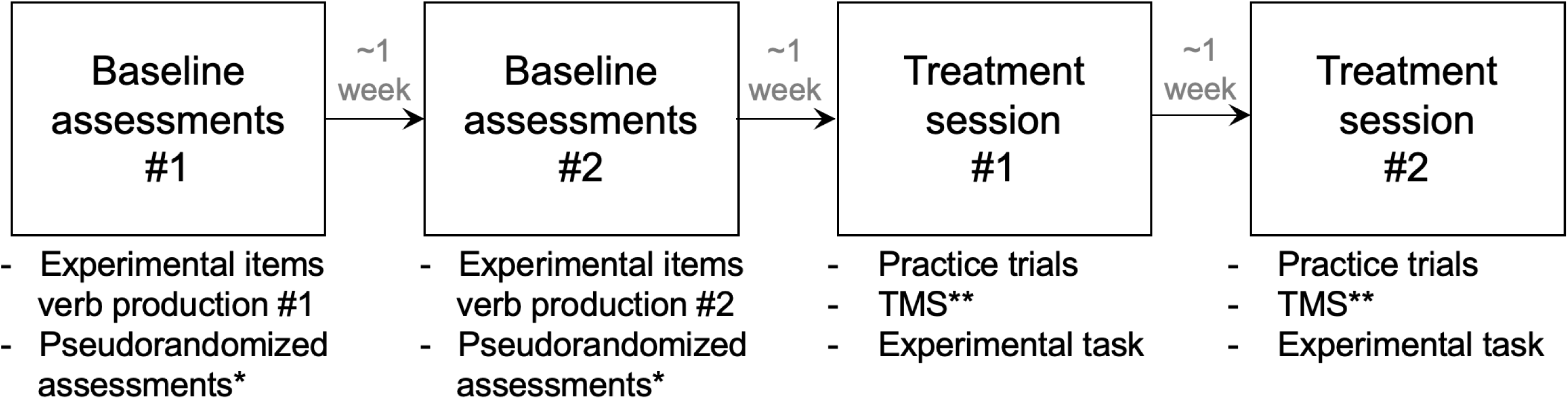
Research session activities. *Notes:* * Pseudorandomized behavioral assessments included: NAVS = Northwestern Assessment of Verbs and Sentences (Cho-Reyes & Thompson, 2012). VNT = Verb Naming Test. VCT = Verb Comprehension Test. ASPT = Argument Structure Production Test. PNT = Philadelphia Naming Test (Roach et al., 1996). KDT = Kissing and Dancing Test (Bak & Hodges, 2003). Event = Event Task (Dresang, Warren, & Dickey, 2019). Meaningful and meaningless gesture imitation and spatial and semantic gesture recognition (Buxbaum et al., 2005, 2014). ** TMS stimulation site was counterbalanced between subjects for order, targeting left pMTG versus vertex. TMS = transcranial magnetic stimulation, in particular intermittent theta burst stimulation. pMTG = posterior middle temporal gyrus.

### Participants

Participants were ten adults with chronic aphasia due to a single left hemisphere ischemic stroke that occurred >1 year prior to participation. See Table 1 for participant demographics. Based on the inclusion and exclusion criteria outlined below, all participants were recruited following their enrollment in R01-DC016800 (PI: H. Branch Coslett). Participants with a history of seizure, psychosis, alcohol and drug abuse, comorbid neurologic deficits other than those engendered by stroke, or any contraindications to TMS were excluded. Inclusion criteria included premorbidly right-handed individuals who were native English speakers and had no lesion overlap with left posterior middle temporal gyrus (pMTG), as defined by Lausanne atlas scale 125 region ID 211 (Hagmann et al., 2008) and validated by a neurologist (See Figure 2). All participants had a research-quality MRI scan and Western Aphasia Battery – Revised (WAB-R; Kertesz, 2006) assessment completed during a separate study in our lab for which they were enrolled less than one year prior to the current study. All participants were in the chronic stage at the time of their MRI and WAB-R completion (> 1-year post-stroke) and medical records reflected no recent changes in language, cognition, or function. All participants were native English speakers, with comprehension abilities adequate to follow instructions (WAB-R comprehension >4), and mild to moderate aphasia based on the WAB-R Aphasia Quotient (Kertesz, 2006; a global measure of language ability). All participants provided informed consent prior to participation, in accordance with the Institutional Review Board of Perelman School of Medicine at the University of Pennsylvania and were paid for their participation.

**Figure 2.**
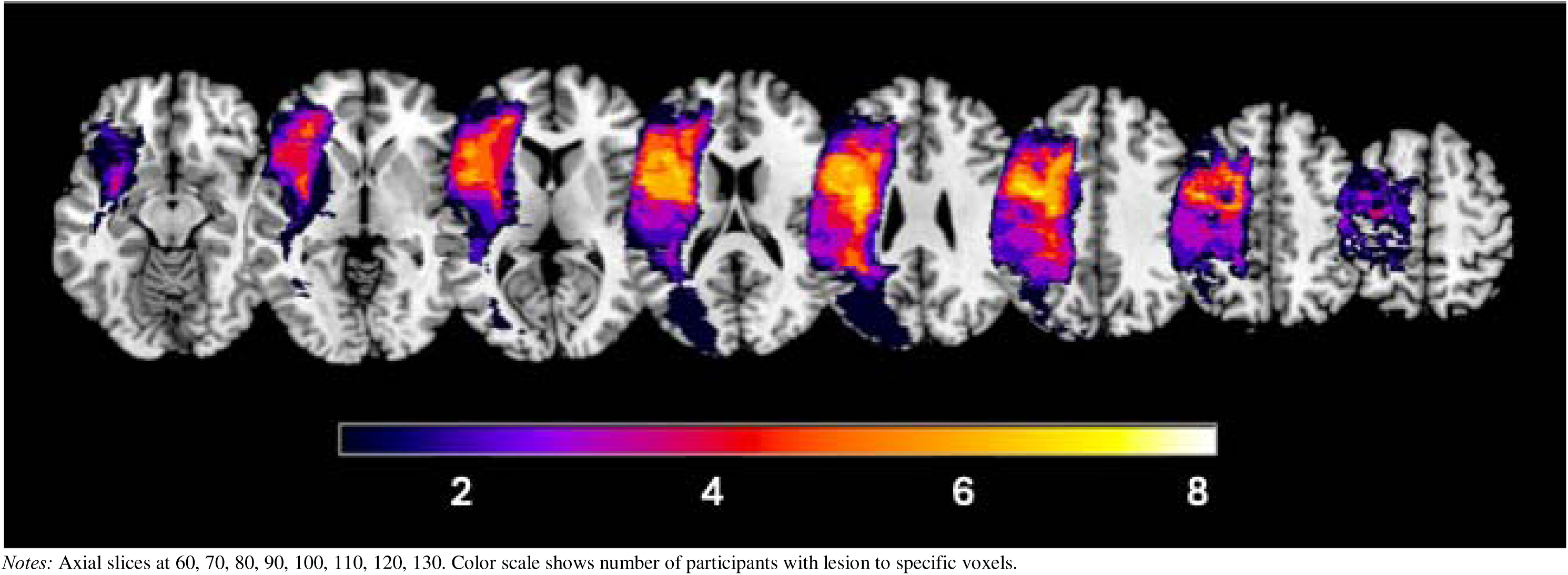
Lesion overlay map. *Notes:* Axial slices at 60, 70, 80, 90, 100, 110, 120, 130. Color scale shows number of participants with lesion to specific voxels.

**Table 1.**
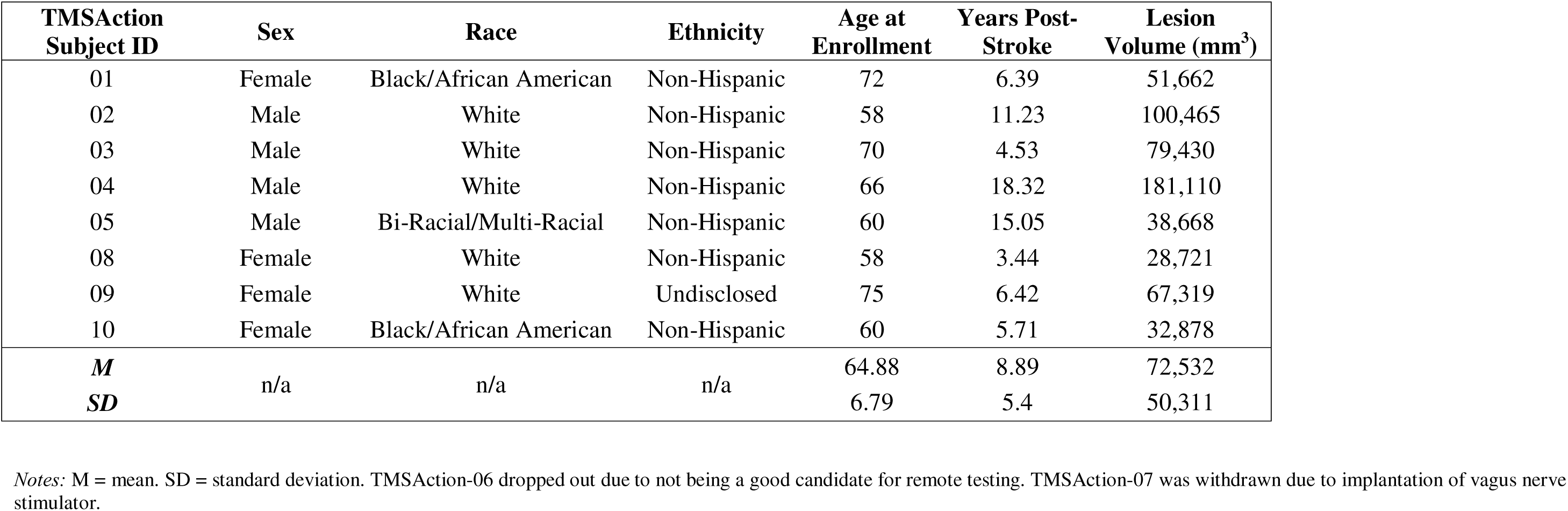
Participant demographic characteristics.

| **TMSAction**<br>**Subject ID** | **Sex** | **Race** | **Ethnicity** | **Age at**<br>**Enrollment** | **Years Post-**<br>**Stroke** | **Lesion**<br>**Volume (mm<sup>3</sup>)** |
| --- | --- | --- | --- | --- | --- | --- |
| 01 | Female | Black/African American | Non-Hispanic | 72 | 6.39 | 51,662 |
| 02 | Male | White | Non-Hispanic | 58 | 11.23 | 100,465 |
| 03 | Male | White | Non-Hispanic | 70 | 4.53 | 79,430 |
| 04 | Male | White | Non-Hispanic | 66 | 18.32 | 181,110 |
| 05 | Male | Bi-Racial/Multi-Racial | Non-Hispanic | 60 | 15.05 | 38,668 |
| 08 | Female | White | Non-Hispanic | 58 | 3.44 | 28,721 |
| 09 | Female | White | Undisclosed | 75 | 6.42 | 67,319 |
| 10 | Female | Black/African American | Non-Hispanic | 60 | 5.71 | 32,878 |
| ***M*** | n/a | n/a | n/a | 64.88 | 8.89 | 72,532 |
| ***SD*** |  |  |  | 6.79 | 5.4 | 50,311 |

*Notes:* M = mean. SD = standard deviation. TMSAction-06 dropped out due to not being a good candidate for remote testing. TMSAction-07 was withdrawn due to implantation of vagus nerve stimulator.

Two participants did not complete the protocol: One completed the baseline phase but declined to participate further, and the second became ineligible after enrollment for medical reasons. In the final set of eight participants, the range of aphasia severity included mild to moderate deficits (aphasia quotient of WAB-R 68.5–84.3). In addition, all subjects scored between 25–85% on each of their baseline assessments of the experimental task, indicating they had sufficient room for improvement and would not show ceiling or floor effects.

### Baseline Assessments

We assessed the following abilities across two baseline sessions: noun production (Philadelphia Naming Test: Roach et al., 1996; from which we derived estimates of semantic versus phonological naming impairments for each subject), verb production (Northwestern Assessment of Verbs & Sentences - Verb Naming Test: Cho-Reyes & Thompson, 2012) sentence production (Northwestern Assessment of Verbs & Sentences - Argument Structure Production Test: Cho-Reyes & Thompson, 2012), verb comprehension (Northwestern Assessment of Verbs & Sentences - Verb Comprehension Test: Cho-Reyes & Thompson, 2012), meaningful and meaningless gesture imitation, gesture recognition with spatial and semantic foils (Buxbaum et al., 2005, 2014), non-linguistic action semantic processing (Kissing and Dancing Test: Bak & Hodges, 2003), and non-linguistic event semantic processing (Event Task: Dresang et al., 2019). We measured verb production on the experimental items at both assessment sessions to provide a more reliable measure of baseline performance. We pseudorandomized the assessment order between subjects. Assessment sessions lasted approximately two hours. See Table 2 for participant performance across tasks.

**Table 2.** Participant performance across tasks.

| Subject ID | WAB-AQ | WAB Classification | experimental task |  |  |  |  |  |  | language |  |  |  | semantic memory |  | gesture |  |  |  |
| --- | --- | --- | --- | --- | --- | --- | --- | --- | --- | --- | --- | --- | --- | --- | --- | --- | --- | --- | --- |
|  |  |  | baseline 1 | baseline 2 | baseline M | neutral × vertex | neutral × pMTG | gesture × vertex | gesture × pMTG | NAVS-VCT | NAVS-VNT | NAVS-ASPT | PNT | Event | KDT | meaningful imitation | meaningless imitation | spatial recognition | semantic recognition |
| 01 | 68.7 | Conduction | 0.375 | 0.281 | 0.328 | 0.40 | 0.484 | 0.69 | 0.556 | 1.00 | 0.500 | 0.313 | 0.480 | 0.879 | 0.558 | 0.879 | 0.667 | 0.70 | 0.80 |
| 02 | 76.4 | Broca's | 0.406 | n/a | 0.406 | 0.60 | 0.714 | 0.759 | 0.621 | 0.955 | 0.682 | 0.813 | 0.815 | 0.930 | 0.865 | 0.800 | 0.700 | 0.50 | 0.90 |
| 03 | 78.3 | Anomic | 0.633 | 0.586 | 0.609 | 0.60 | 0.714 | 0.69 | 0.793 | 1.00 | 0.818 | 0.406 | 0.682 | 0.949 | 0.712 | 0.700 | 0.767 | 0.80 | 0.90 |
| 04 | 77.6 | Conduction | 0.523 | 0.555 | 0.539 | 0.793 | 0.724 | 0.657 | 0.771 | 1.00 | 0.364 | 0.750 | 0.759 | 0.930 | 0.904 | 0.967 | 0.767 | 0.80 | 1.00 |
| 05 | 83.8 | Anomic | 0.430 | n/a | 0.430 | 0.667 | 0.75 | 0.735 | 0.694 | 1.00 | 1.00 | 1.00 | 0.817 | 0.906 | 0.808 | 0.733 | 0.567 | 0.60 | 0.90 |
| 08 | 84.3 | Anomic | 0.805 | 0.820 | 0.813 | 0.781 | 0.938 | 0.906 | 0.906 | 1.00 | 0.955 | 1.00 | 0.920 | 0.980 | 0.962 | 0.933 | 0.933 | 0.70 | 0.90 |
| 09 | 77.1 | Transcortical Motor | 0.422 | 0.405 | 0.413 | 0.314 | 0.686 | 0.621 | 0.448 | 0.773 | 0.773 | 0.656 | 0.723 | 0.809 | 0.827 | 0.800 | 0.767 | 0.70 | 0.80 |
| 10 | 68.5 | Anomic | 0.258 | 0.313 | 0.285 | 0.40 | 0.343 | 0.448 | 0.379 | 0.682 | 0.409 | 0.156 | 0.366 | 0.633 | 0.577 | 0.433 | 0.400 | 0.50 | 0.50 |
| **M** | **76.84** | **n/a** | **0.48** | **0.49** | **0.48** | **0.57** | **0.67** | **0.69** | **0.65** | **0.93** | **0.69** | **0.64** | **0.70** | **0.88** | **0.78** | **0.77** | **0.70** | **0.64** | **0.83** |
| **SD** | **5.89** | **n/a** | **0.17** | **0.20** | **0.17** | **0.18** | **0.18** | **0.13** | **0.18** | **0.13** | **0.24** | **0.32** | **0.18** | **0.11** | **0.15** | **0.18** | **0.17** | **0.11** | **0.16** |

*Notes:* M = mean. SD = standard deviation. WAB-AQ = Western Aphasia Battery – Aphasia Quotient (Kertesz, 2006). TMS = transcranial magnetic stimulation, in particular intermittent theta burst stimulation. pMTG = posterior middle temporal gyrus. NAVS = Northwestern Assessment of Verbs and Sentences (Cho-Reyes & Thompson, 2012). VNT = Verb Naming Test. VCT = Verb Comprehension Test. ASPT = Argument Structure Production Test. PNT = Philadelphia Naming Test (Roach et al., 1996). KDT = Kissing and Dancing Test (Bak & Hodges, 2003). Event = Event Task (Dresang, Warren, & Dickey, 2019). Gesture imitation and recognition tasks evaluated meaningful and meaningless gesture production and semantic and spatial gesture recognition components (Buxbaum et al., 2005, 2014).

### Experimental Task

All subjects underwent two experiment sessions where they completed the same behavioral task during cross-over sessions of iTBS to left pMTG or vertex control site. We pseudorandomized the order of stimulation site and scheduled iTBS sessions at least one week apart. The behavioral task, adapted from Murteira & Nickels (2020), required participants to use a complete sentence to describe action pictures following the presentation of congruent or neutral gesture primes. Stimuli included 128 action pictures, 128 congruent gesture videos, and a set of four neutral gesture videos. All pictures were 300 x 300 pixel, black-and-white line drawings, retrieved from a variety of standardized action picture collections whenever possible (Druks & Masterson, 2000; Murteira et al., 2018; Murteira & Nickels, 2020; Szekely et al., 2004). Videos were retrieved from previous research datasets and opensource American Sign Language videos (Murteira & Nickels, 2020; Ortega et al., 2019). All videos were edited to contain no audio or movement outside of the target gesture (e.g., mouth movements, facial expressions, grooming movements), consistent centered framing of the torso, and full completion of the gesture lasting 900 ms.

All pictures and video clips were normed for name agreement, familiarity, and ratings for how strongly each gesture mapped onto a single lexical form. Normative surveys were administered online via *Amazon Mechanical Turk* (n.d.), a crowdsourcing platform commonly used for behavioral research. All survey participants self-reported as being from the United States, aged 18 to 79, with English as their primary language. They reported no colorblindness or other vision issues precluding participation. To ensure survey data validity, several quality control procedures were implemented. Only fully completed surveys were included, and responses were evaluated for signs of insufficient engagement (e.g., abnormally short completion times) and irregular response patterns. Analyses were conducted on responses that satisfied these quality standards. Action picture-name agreement averaged 71% (SD = 22%, N = 26), while gesture-name agreement was 66% on average (SD = 30%, N = 25). Of note, these measures are specific to a single target name, with the intent to evaluate the degree of inter-individual agreement of lexical mapping, and thus synonyms or otherwise acceptable alternatives were not counted as agreement. In addition, these norms informed acceptable alternative responses for the experimental task (see *Scoring Procedures*). Participants rated action picture familiarity on a 1–5 scale (M = 3.84, SD = 1.05, N = 25). The gesture-name fit or appropriateness was also rated on a 1–5 scale (M = 4.15, SD = 0.83, N = 27). These results were largely consistent with previous studies (Murteira & Nickels, 2020), and therefore all normed stimuli were retained.

We created two sets of items, such that half the items presented were preceded by congruent gesture primes (Set A) and half were preceded by neutral primes; that is, an actor standing with no arm/hand movement (Set B). In the second experiment session, the items were preceded by the opposite prime. Thus, in each experimental session, participants named each target picture only once, but by the end of the two sessions, all participants had encountered each action picture in both gesture conditions. Sets A-B were balanced for psycholinguistic properties of the target verbs. To ensure balance across condition sets, one-way ANOVAs were conducted on several verb-related variables. No significant differences were found between condition sets for action familiarity, F(1, 126) = 0.08, p = .772; name agreement, F(1, 126) = 0.29, p = .591; number of phonemes, F(1, 126) = 0.03, p = .874; instrumentality, F(1, 126) = 0.03, p = .856; transitivity, F(1, 126) = 0.04, p = .851; lexical frequency, F(1, 126) = 0.01, p = .932; or age of acquisition, F(1, 126) = 0.03, p = .859. Moreover, previous research indicates that these factors are unlikely to influence response to verb naming facilitation following gesture observation (Murteira & Nickels, 2020).

The experimental task was presented on a computer screen. Each trial included: (1) a fixation cross appearing in the middle of the screen for 1000 milliseconds (ms); (2) a pantomime gesture video clip for 900 ms; (3) a fixation cross appearing in the middle of the screen for 100 ms; and (4) an action picture appearing in the middle of the screen for 5000 ms. Prior to the task, we told participants that brief video clips of gestures and pictures of actions would appear on the screen. We then asked participants to name the pictures of actions using a complete sentence with a single verb gerund phrase (e.g., *he is drinking*), and we instructed them not to respond to the video clips. Verbal and written instructions were provided, and six practice trials were performed to ensure adequate comprehension of the task. Presentation order was pseudorandomized across participants. Specifically, each participant completed 128 trials across four experimental sessions, resulting in a total of 512 observations per participant and 4,096 observations overall.

### Scoring Procedures

Phrases containing the target verb (e.g., “drinking” or “The boy is drinking water.”) were scored as correct. A maximum of one prompt was provided per trial. Following the procedure by Murteira and colleagues (2020), correct responses were counted if the participant produced: 1) the target verb with no phonological error, 2) the target verb with a phonological error but sharing >50% of phonemes with the stem of the target verb (e.g., /dink ŋ/ for *drinking*), or 3) a self-correction by the subject. Since participants did not receive feedback on their responses, any form of the target verb was accepted (e.g., *drink*, *drinks*, *drank*). and synonyms were accepted based on the name agreement norms, Merriam-Webster dictionary definitions, and consensus judgments of two independent raters (e.g., *scratch, itch*) who transcribed and coded all data. The raters demonstrated inter-rater reliability for 75% of the items, resulting in an observed agreement of 98.11%. Cohen’s kappa for inter-rater agreement was 0.936, indicating a very high level of agreement beyond chance. All discrepancies between the raters were discussed and resolved through consensus judgments.

### Neuroimaging Procedures

All participants underwent high-resolution magnetic resonance imaging (MRI) brain scans on a Siemens 3T Prisma prior to the baseline assessment sessions. The MRI scan protocol included high-resolution T1-weighted scans (1 × 1 × 1 mm voxels), high-resolution diffusion imaging (HARDI), fluid attenuated inversion recovery (FLAIR), and pseudo-CASL perfusion imaging. To characterize the extent and location of lesions, we followed lesion tracing procedures outlined in Schnur et al. (2009) and employed in numerous other studies (Dresang et al., 2025; Schwartz et al., 2011, 2012).

Finally, to compute the disconnectivity metrics, we mapped the lesion of each participant onto tractography reconstructions of white matter pathways obtained from a group of healthy controls (Rojkova et al., 2016). We quantified the severity of the disconnection by measuring the probability that the tract was disconnected and the proportion of disconnection (Thiebaut de Schotten et al., 2014) using Tractotron software as part of the BCBtoolkit (http://www.brainconnectivitybehaviour.eu). The small sample size precludes analyses of the relationship of the neuroanatomical data to the behavioral indices, but we include them as points of potential interest for future research (see *Supplementary Materials*).

### TMS Procedures

To set stimulation sites for neuronavigation, we loaded each participant’s T1 MRI scan into Brainsight, a neuronavigation software that rendered curvilinear models of each participant’s brain (Rogue Research, 2025). We mapped each brain model to MNI space (Montreal Neurological Institute; Mazziotta et al., 1995a, 1995b) and localized the primary stimulation target, the left pMTG, for each subject by inputting MNI coordinates (−45, −69, 5). We selected the vertex control site, defined as the scalp location corresponding to electrode site Cz in the international 10–20 EEG system. Anatomically, this lies on the midline cortical point of the sagittal suture, between the precentral and postcentral gyri, at the highest point of the skull. Brainsight software optimized coil positioning and trajectories using the rendered scalp surface model generated for each participant.

During the experimental tasks, iTBS was applied to the left pMTG or the vertex control site. Standard iTBS parameters were used, with stimulation applied at 80% of active motor threshold (AMT; Huang et al., 2005). The iTBS protocol involves bursts containing three pulses at 50 Hz repeated at 200-ms intervals for two seconds (i.e., at 5 Hz). A two-second train of iTBS was repeated every 10 seconds for a total of 190 seconds and 600 pulses. See Figure 3. Each subject’s AMT was determined at the start of each session.

**Figure 3.**
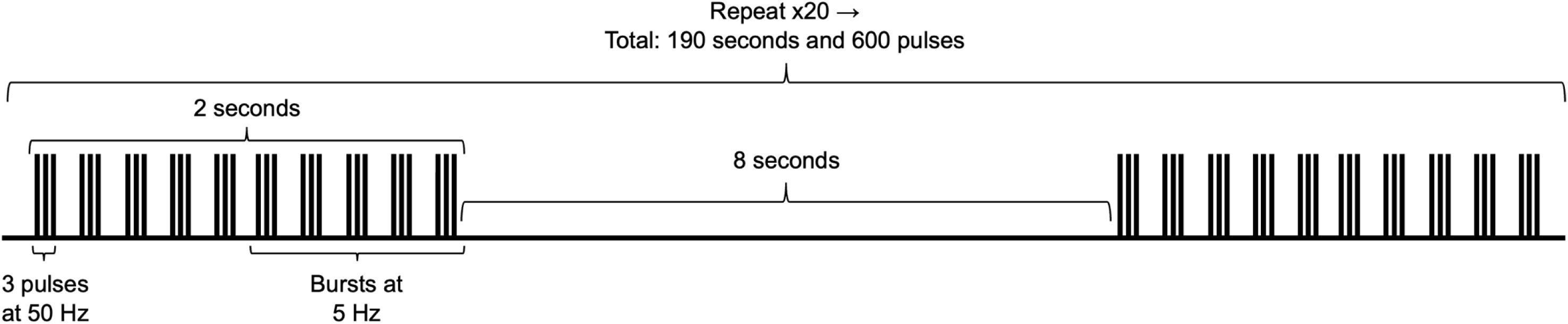
Intermittent theta burst stimulation (iTBS) protocol design.

### Statistical Analysis

This pilot study was designed to establish feasibility of a novel intervention design. A priori power calculations were based on Murteira and Nickels (2020), who reported a moderate effect size (Cohen’s *d* = 0.61) for improved verb retrieval following the observation of congruent versus neutral gestures in ten individuals with aphasia. Using this effect size and the *pwr* package in R (Champeley, 2020), we calculated that a sample size of 23 participants would be required to achieve 80% power. Our original aim was to recruit 30 participants with aphasia to ensure adequate power for the behavioral and TMS conditions, with our initial study design including two planned TMS comparisons (pMTG vs vertex and anterior IFG vs vertex). However, due to recruitment and feasibility constraints during the COVID-19 pandemic, the final sample included 8 participants, and we thus decided to focus on a single TMS contrast of pMTG vs vertex. This reduction in the number of planned comparisons eliminates the need for multiple comparison corrections between stimulation sites, thereby simplifying interpretation and preserving statistical power for the remaining contrast. This sample size is consistent with other pilot TMS studies in clinical populations and provides valuable initial data to evaluate the experimental protocol, estimate variability, and inform the design, hypotheses, and power analysis of future, fully powered studies. Additionally, there was a large number of within-subject trials, which supports reliable estimation of fixed effects in the generalized linear mixed-effects model, particularly for the within-subject interaction of prime × TMS.

We implemented Bayesian generalized linear mixed-effects models using Stan with the brms package in the R statistical software version (Bürkner, 2017; Carpenter et al., 2017). This modeling approach provides more stable uncertainty quantification under small-N multilevel conditions by quantifying the probability and uncertainty of the prime × TMS interaction under hierarchical constraints. We analyzed the effects of gesture priming (gesture vs. neutral), stimulation site (pMTG vs. vertex), and their interaction on verb retrieval accuracy. We controlled for the effects of TMS session order and baseline accuracy (averaged and centered), and we included random intercepts for subjects (N = 8) and items (N = 128) to account for individual and lexical variability.

We modeled trial-level accuracy using a Bayesian logistic regression with a Bernoulli likelihood and logit link. Following common recommendations for weakly informed priors in logistic models (e.g., Gelman et al., 2008), regression coefficients were assigned normal priors centered at zero with a standard deviation of 1.5. Posterior distributions were estimated using Hamiltonian Monte Carlo as implemented in Stan (Carpenter et al., 2017). We ran four Markov Chain Monte Carlo (MCMC) chains, with 4,000 iterations per chain, including 2,000 warm-up iterations, resulting in 8,000 retained posterior draws. Consistent with recommended Bayesian workflow practices (e.g., Gelman et al., 2020), posterior summaries included means, standard errors, 95% credible intervals (CI). To quantify evidence for directional effects, we also report the proportion of posterior samples supporting a directional effect (e.g., P(β > 0) or P(β < 0)).

Model convergence was assessed using the Gelman-Rubin potential scale reduction statistic (R□), the number of effective samples, and visual inspection of trace plots (e.g., Gelman & Rubin, 1992). Model fit was examined using posterior predictive checks (e.g., Gelman et al., 1996). The primary hypothesis was assessed by testing the interaction between gesture prime and TMS site, controlling for session order and baseline performance. Pairwise comparisons were then conducted using estimated marginal means (*emmeans* package; Lenth et al., 2022), with Bonferroni correction applied to control for multiple comparisons.

## RESULTS

All model parameters showed satisfactory convergence. The Gelman–Rubin statistic (R□) was 1.00 for all parameters, indicating that independent chains converged to the same posterior distribution (e.g., Gelman & Rubin, 1992). Effective sample sizes were sufficiently large, and no divergent transitions occurred, suggesting that the Hamiltonian Monte Carlo sampler explored the posterior distribution reliably (e.g., Betancourt, 2017). We found that the gesture priming × TMS interaction was a reliable predictor of verb retrieval accuracy (β = −0.64, SE = 0.32, 95% CI: [−1.26, −0.02]), with 97.9% of posterior samples below zero, indicating strong evidence for a negative interaction. See Figure 4 for the predicted probability of verb accuracy interaction effect. Surprisingly, this corresponded to a 47% reduction in the odds of correct naming (OR = 0.53, 95% CI: [0.28, 0.98]). By contrast, the main effects of priming (β = 0.11, SE = 0.24, 95% CI: [−0.36, 0.59]; OR = 1.12, 67.3% posterior > 0) and TMS (β = 0.13, SE = 0.23, 95% CI: [−0.25, 0.36]; OR = 1.14, 71.6% posterior > 0) were weak and less conclusive. The estimated random intercepts for subjects (SD = 0.69, 95% CI [0.27, 1.39]) and items (SD = 0.91, 95% CI [0.66, 1.18]) indicate that there was substantial variability in naming accuracy. Table 3 shows posterior estimates for all fixed effects.

**Figure 4.**
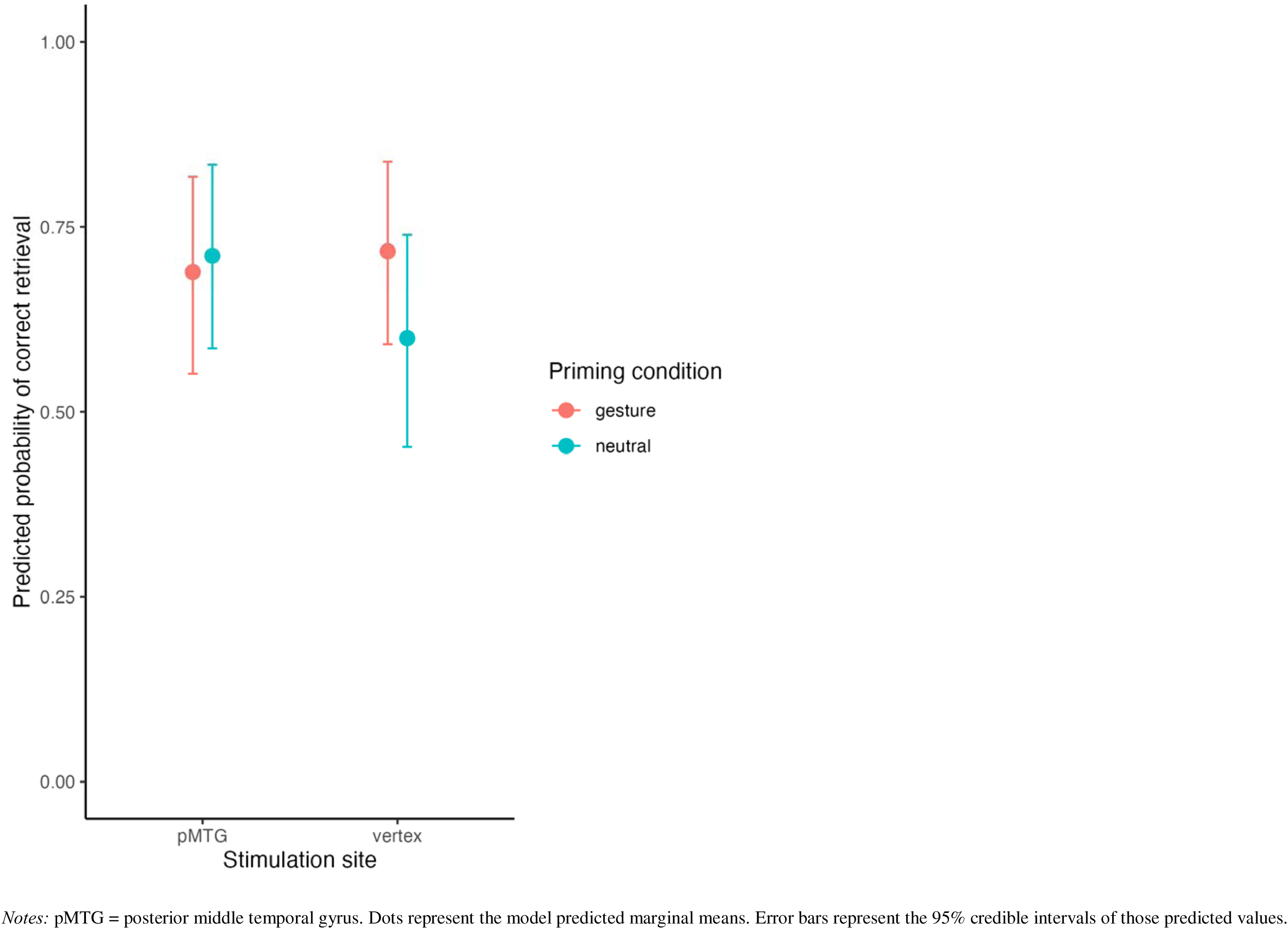
Predicted probability of verb accuracy model interaction effect. *Notes:* pMTG = posterior middle temporal gyrus. Dots represent the model predicted marginal means. Error bars represent the 95% credible intervals of those predicted values.

**Table 3.** Posterior estimates for all fixed effects.

| Predictor | Estimate | Est. Error | 95% CI |
| --- | --- | --- | --- |
| intercept | 0.76 | 0.32 | 0.11, 1.41 |
| priming | 0.11 | 0.24 | −0.36, 0.59 |
| TMS | 0.13 | 0.23 | −0.32, 0.57 |
| order | 0.06 | 0.15 | −0.25, 0.36 |
| baseline | 1.50 | 1.05 | −0.59, 3.48 |
| priming : TMS | −0.64 | 0.32 | −1.26, −0.02 |

Notes: Intercept represents neutral + vertex condition. Order reflects the counterbalanced order of TMS site. Baseline performance on the naming task was averaged across two independent observations and centered.

Emmeans on the model revealed the predicted probabilities of verb retrieval for each priming × TMS condition combination (Table 4). Under vertex stimulation, gesture priming increased predicted naming accuracy by 12 percentage points (0.72 vs. 0.60). In the absence of gesture priming, pMTG stimulation likewise improved performance relative to vertex stimulation by 11 percentage points (0.71 vs. 0.60). Critically, however, these benefits were not additive: when gesture priming and pMTG stimulation were combined, performance did not exceed baseline levels (0.69). This pattern produced the reliable negative interaction observed in the model, indicating that left pMTG stimulation attenuated the facilitative effect of gesture priming and/or gesture attenuated the facilitative effect of pMTG stimulation. In contrast, the individual main effects of priming and TMS were weak and unreliable.

**Table 4.** Marginal effects predicted probabilities of correct verb retrieval.

| Priming Condition | TMS Condition | Predicted Probability | 95% CI |
| --- | --- | --- | --- |
| neutral | vertex | 0.60 | 0.45, 0.74 |
| neutral | pMTG | 0.71 | 0.59, 0.83 |
| gesture | vertex | 0.72 | 0.59, 0.84 |
| gesture | pMTG | 0.69 | 0.55, 0.82 |

Notes: pMTG = left posterior middle temporal gyrus. CI = credible interval.

## DISCUSSION

Strengthening the semantic associations between verbs and other entities associated with them (e.g., people, locations, and gestures) has been a beneficial treatment approach for participants with a range of aphasia types, severities, and culturolinguistic backgrounds (e.g., Edmonds, 2016). However, standard language therapies, including verb-strengthening treatments, often demonstrate limited and inconsistent efficacy for individuals with chronic aphasia (e.g., Peitz et al., 2024). This preliminary investigation serves as the first pilot study to examine whether combining gestural and TMS activation of action semantics can facilitate verb retrieval in adults with chronic aphasia.

Using a Bayesian generalized linear mixed-effects model, we observed evidence for a negative interaction between gesture priming and stimulation site. Contrary to our predictions, the combination of gesture priming with pMTG stimulation did not yield additional benefits over either intervention alone; rather, pMTG stimulation attenuated the priming advantage observed under vertex stimulation, and conversely, gesture attenuated the advantage we observed with pMTG stimulation alone. This non-additive pattern is reflected in the predicted probabilities, where performance in the combined condition (gesture + pMTG) did not exceed that observed in either intervention alone. The posterior estimate for the interaction was negative, with a 95% credible interval that narrowly excluded zero, indicating evidence for an interaction, whereas the main effects of priming and stimulation were small and associated with considerable uncertainty. Using a Bayesian framework permits a graded interpretation of this non-additive pattern, indicating that the data are substantially more compatible with a negative interaction than with a null effect.

The observed pattern of gesture priming under vertex stimulation was numerically consistent with prior work demonstrating that meaningful gestures can facilitate lexical access, potentially by strengthening conceptual activation and promoting spreading activation to the lexical word form (Kalénine & Buxbaum, 2016; Lebkuecher et al., 2026; Murteira et al., 2019; Murteira & Nickels, 2020; Rose, 2006). However, in the present data, the main effect of gesture priming was associated with substantial uncertainty and therefore does not provide strong standalone evidence for facilitation. Likewise, although there was a numeric facilitative effect of pMTG stimulation in the neutral condition, the estimated main effect of stimulation was uncertain. This pattern is nonetheless broadly consistent with prior evidence implicating left pMTG in lexical-semantic processing, action concept representation, and verb retrieval (Andric & Small, 2012; Dresang et al., 2023; Kabulska et al., 2024; Peelen et al., 2012; Straube et al., 2012; Tarhan et al., 2015; Wurm & Caramazza, 2019). Together, these findings support the view that both gesture observation and neuromodulation of pMTG can independently enhance access to action semantics and facilitate verb retrieval in a sentence production paradigm. The current findings should be interpreted in the context of a small pilot sample, which limits the precision of the estimated effects. Within this context, the observed directional trends remain informative, particularly insofar as they align with prior theoretical and empirical work.

Importantly, however, the interaction challenges both additive and positively interactive models in which gestures and pMTG stimulation together are particularly facilitative. One plausible interpretation of the pattern of results is based on the model that gesture observation and pMTG stimulation facilitate the same action semantic representations, albeit via different input mechanisms (visual versus neuromodulatory input). If pMTG stimulation upregulates this network, gesture-based conceptual activation may provide reduced additional benefit when both are combined, resulting in a sub-additive pattern that is consistent with overlapping or partially competing mechanisms. One possible account is that simultaneous upregulation from both sources increases activation within the semantic system in a way that also amplifies competition among candidate representations. Within spreading activation frameworks, lexical retrieval depends not only on activation of target representations but also on the resolution of competition, which can place demands on selection and control mechanisms (e.g., Dell, 1986; Jefferies et al., 2006; Thompson-Schill et al., 1997). Under such conditions, additional input may not translate into improved performance if it disproportionately increases competition or selection demands, resulting in the observed plateau in behavior. This interpretation is consistent with neurocognitive accounts of semantic processing that emphasize the interaction between representational activation and control processes across distributed cortical networks (Lambon Ralph et al., 2017; Patterson et al., 2007; Rogers et al., 2015). From this perspective, the present findings suggest that combined gesture and pMTG stimulation may not yield additive benefits because both manipulations engage overlapping components of the semantic system, with performance potentially constrained by the efficiency of selection rather than the absolute level of activation.

Substantial variability was observed across participants and items, as reflected in the magnitude of the random intercept estimates. Although we cannot directly attribute this variability to specific neuroanatomical or behavioral predictors in the present dataset due to the small sample size, its presence underscores the heterogeneity observed in chronic aphasia and neuromodulation response. In particular, our data suggests considerable inter-individual differences not only in effect size but also in the direction of TMS-related change, with some individuals showing excitatory responses and others inhibitory responses to pMTG stimulation. Such variability is consistent with prior TMS literature and may reflect differences in lesion distribution, residual structural connectivity, neuroplastic potential, or baseline network efficiency (e.g., Dresang et al., 2022; Parchure et al., 2022; Shah-Basak et al., 2020). Furthermore, substantial inter individual variability in TBS aftereffects has been documented even in neurologically healthy samples, with responsiveness depending on baseline cortical excitability and other neurophysiological factors (Corp et al., 2020; Wischnewski & Schutter, 2015). It will be of interest to examine potential behavioral and neuroanatomic predictors of response in larger future samples.

These findings have both theoretical and clinical implications. From a theoretical perspective, the results support models in which gesture observation cues and left pMTG stimulation modulate a shared semantic repository rather than operating via independent pathways. Under classic additive-factor logic (Sternberg, 1969, 1998), factors that influence distinct processing stages are expected to produce additive effects on performance, whereas deviations from additivity (i.e., interactions) suggest a common locus of influence. In the present data, the non-additive pattern observed when gesture cues and stimulation are combined is therefore more consistent with both conditions engaging overlapping semantic representations or retrieval systems. This interpretation aligns with evidence that gesture and language recruit overlapping neural and representational systems for semantic processing, particularly within left temporal cortex (Andric & Small, 2012; Willems et al., 2007).

From a clinical standpoint, gesture cues appear to provide a reliable and low-barrier facilitative strategy in the absence of neuromodulation, consistent with prior evidence (e.g., Murteira & Nickels, 2020; Rose, 2006). In contrast, the clinical utility of left pMTG stimulation for supporting verb retrieval remains uncertain, given its context-dependent effects and inter-individual variability. More broadly, the present findings raise the possibility that combining neuromodulation with behavioral interventions may not uniformly yield additive—or even synergistic—benefits. Although pairing stimulation with therapy is often assumed to maximize treatment gains, our results suggest that outcomes may depend on the specific linguistic function targeted, its neural instantiation, and the degree to which the behavioral and neurostimulatroy manipulations converge on shared neurocognitive mechanisms. In some cases, concurrent interventions may compete for or saturate the same underlying representations, thereby attenuating combined effects. Future work should therefore focus on identifying predictors of TMS responsiveness and optimizing stimulation–behavior pairings, potentially through individualized functional imaging or connectivity-based targeting approaches.

There were several limitations to this pilot study, which was conducted in a convenience sample during the COVID-19 pandemic. While fixed effects and group-level interactions were estimated from over 4,000 trial-level observations, the participant sample was small, limiting statistical power, generalizability, and the stability of individual-level estimates. The Bayesian model indicates moderate evidence for an interaction, but uncertainty remains regarding its precise magnitude. Furthermore, the observed heterogeneity suggests that larger samples will be essential to disentangle systematic predictors of treatment response from sampling variability. An additional limitation concerns the normative properties of the stimulus set, particularly low gesture–name agreement. Although normative data were collected to characterize stimulus–label mappings, variability in agreement may have attenuated priming effects by weakening the consistency with which gestures cued the intended verbs. However, the presence of an overall gesture cueing effect suggests that this issue did not nullify the effectiveness of the manipulation. Accordingly, as a feasibility pilot study conducted under constrained conditions, the present findings should be interpreted as preliminary.

Despite these limitations, this study provides the first evidence examining the combined and independent effects of gesture priming and pMTG iTBS on verb retrieval in chronic aphasia. The results suggest that multimodal activation of action semantics does not necessarily produce additive or interactive gains but instead operates in a context-dependent manner. By quantifying evidence on a probabilistic scale and highlighting substantial individual variability, this work establishes an empirical foundation for future studies aimed at optimizing neuromodulatory and behavioral interventions for verb and sentence production deficits.

## CONCLUSION

Our results support the hypothesis that non-invasive brain stimulation and gesture observation can modulate verb retrieval in aphasia. While the precise neurocognitive mechanisms underlying their effects remain to be clarified, these findings align with a growing body of evidence highlighting the facilitative role of gesture cues and non-invasive neurostimulation in language production. However, our results caution against assuming that neurostimulation will enhance outcomes of gesture priming without additional testing in larger samples. Future research should continue investigating the dynamics between neurostimulation, gesture processing, and lexical retrieval, with the goal of optimizing multimodal communication strategies in aphasia rehabilitation.

## Acknowledgments

Felipe Rosero Castro, Rand Williamson, Denise Harvey, Isabella Strauss, Hadeel Manimaran, Apoorva Kelkar, Olu Faseyitan, H. Branch Colsett.

## SUPPLEMENTARY MATERIALS

### Supplementary Tables

Disconnectome metrics. For each participant’s lesion, disconnectome metrics were computed using Tractotron software in the BCBtoolkit (Thiebaut de Schotten et al., 2014) to indicate tract-specific probability and proportion of disconnection on the MNI152 atlas.

**Table S1.** Probability of disconnection per tract and participant.

| Subject ID | Left Arcuate Fasciculus Long Segment | Left Arcuate Fasciculus Posterior Segment | Left Frontal Aslant Tract | Left Frontal Inferior Longitudinal Fasciculus | Left Frontal Superior Longitudinal Fasciculus | Left Inferior Fronto Occipital Fasciculus | Left Inferior Longitudinal Fasciculus | Left Superior Longitudinal Fasciculus I | Left Superior Longitudinal Fasciculus II | Left Superior Longitudinal Fasciculus III | Left Uncinate Fasciculus |
| --- | --- | --- | --- | --- | --- | --- | --- | --- | --- | --- | --- |
| 1 | 1.00 | 1.00 | 1.00 | 0.98 | 1.00 | 1.00 | 0.79 | 1.00 | 1.00 | 1.00 | 0.78 |
| 2 | 1.00 | 1.00 | 1.00 | 1.00 | 1.00 | 1.00 | 0.88 | 1.00 | 1.00 | 1.00 | 0.96 |
| 3 | 1.00 | 0.96 | 1.00 | 1.00 | 1.00 | 1.00 | 0.32 | 1.00 | 1.00 | 1.00 | 1.00 |
| 4 | 1.00 | 1.00 | 1.00 | 1.00 | 1.00 | 1.00 | 1.00 | 1.00 | 1.00 | 1.00 | 1.00 |
| 5 | 1.00 | 0.00 | 1.00 | 1.00 | 1.00 | 0.00 | 1.00 | 1.00 | 1.00 | 1.00 | 0.96 |
| 8 | 0.74 | 0.00 | 1.00 | 1.00 | 1.00 | 0.76 | 0.00 | 0.97 | 1.00 | 0.98 | 0.88 |
| 9 | 1.00 | 0.00 | 1.00 | 1.00 | 1.00 | 1.00 | 0.23 | 1.00 | 1.00 | 1.00 | 1.00 |
| 10 | 1.00 | 1.00 | 1.00 | 0.84 | 0.80 | 1.00 | 1.00 | 1.00 | 1.00 | 1.00 | 1.00 |

**Table S2.**
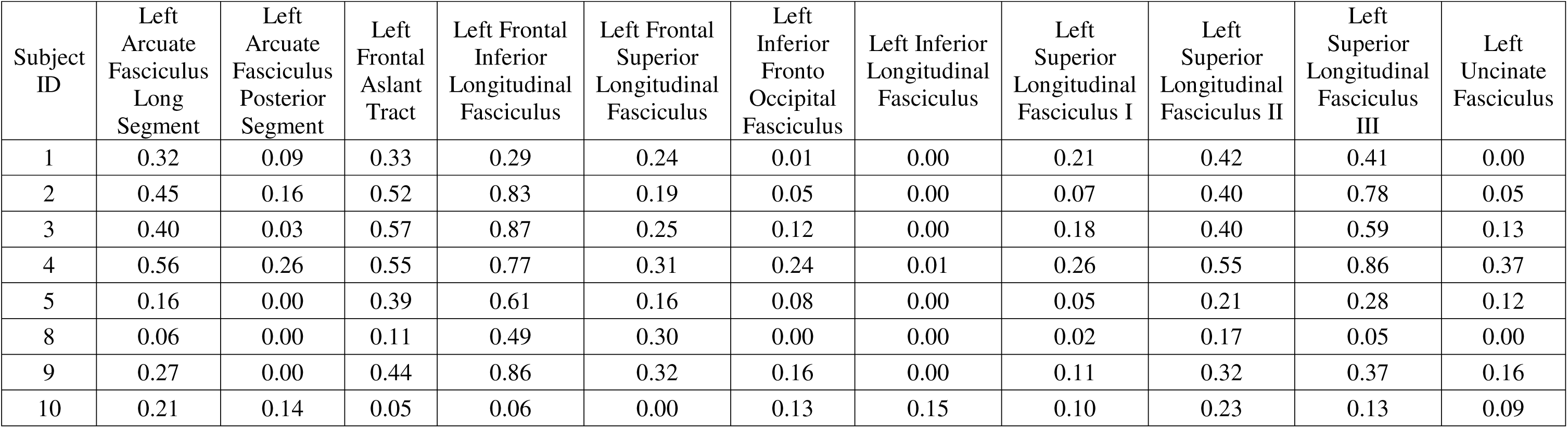
Proportion of disconnection per tract and participant.

### Supplementary Figures

Disconnectome maps. For each participant’s lesion, disconnectome maps were computed using Tractotron software in the BCBtoolkit (Thiebaut de Schotten et al., 2014) to indicate the voxel-wise probability of disconnection on the MNI152 atlas.

**Figure S1.**
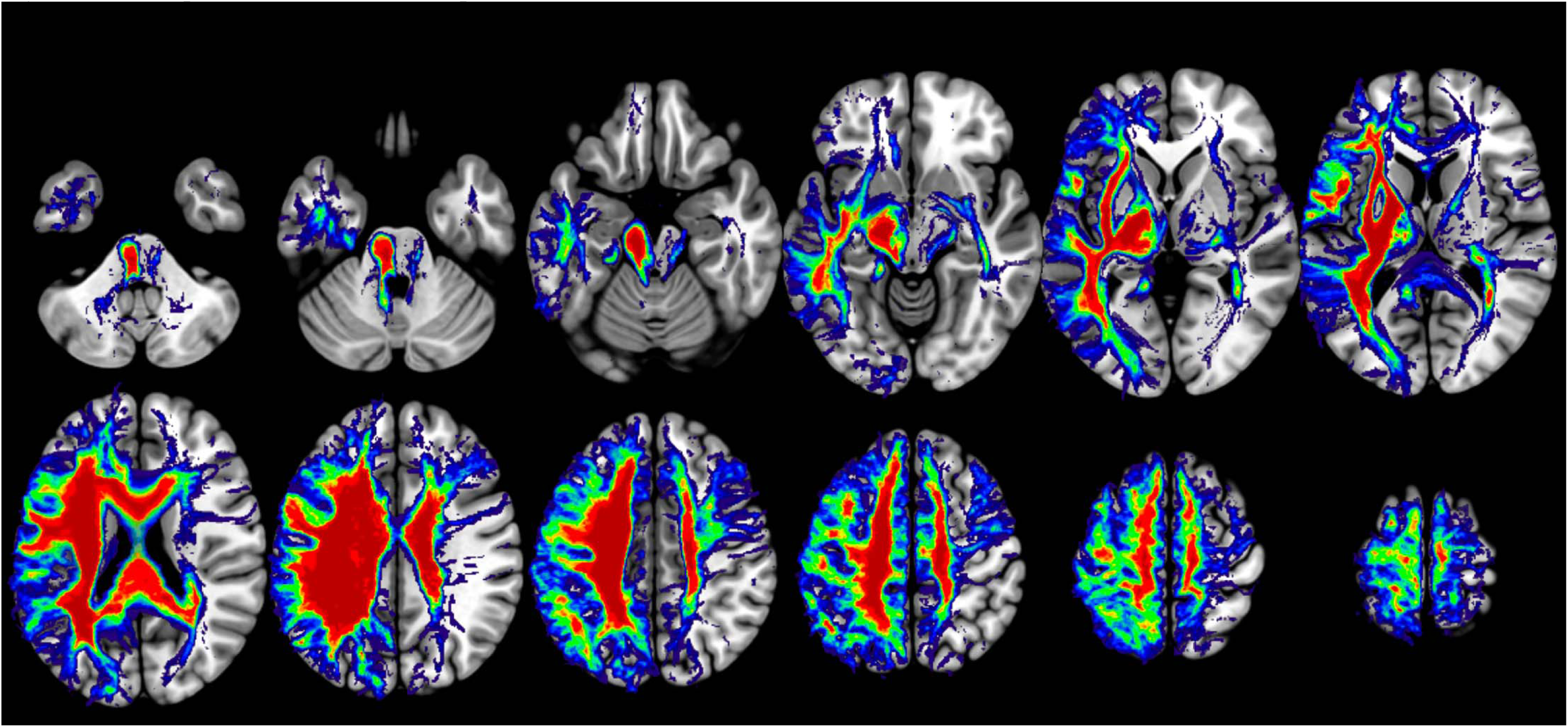
Participant 01 disconnectome map.

**Figure S2.**
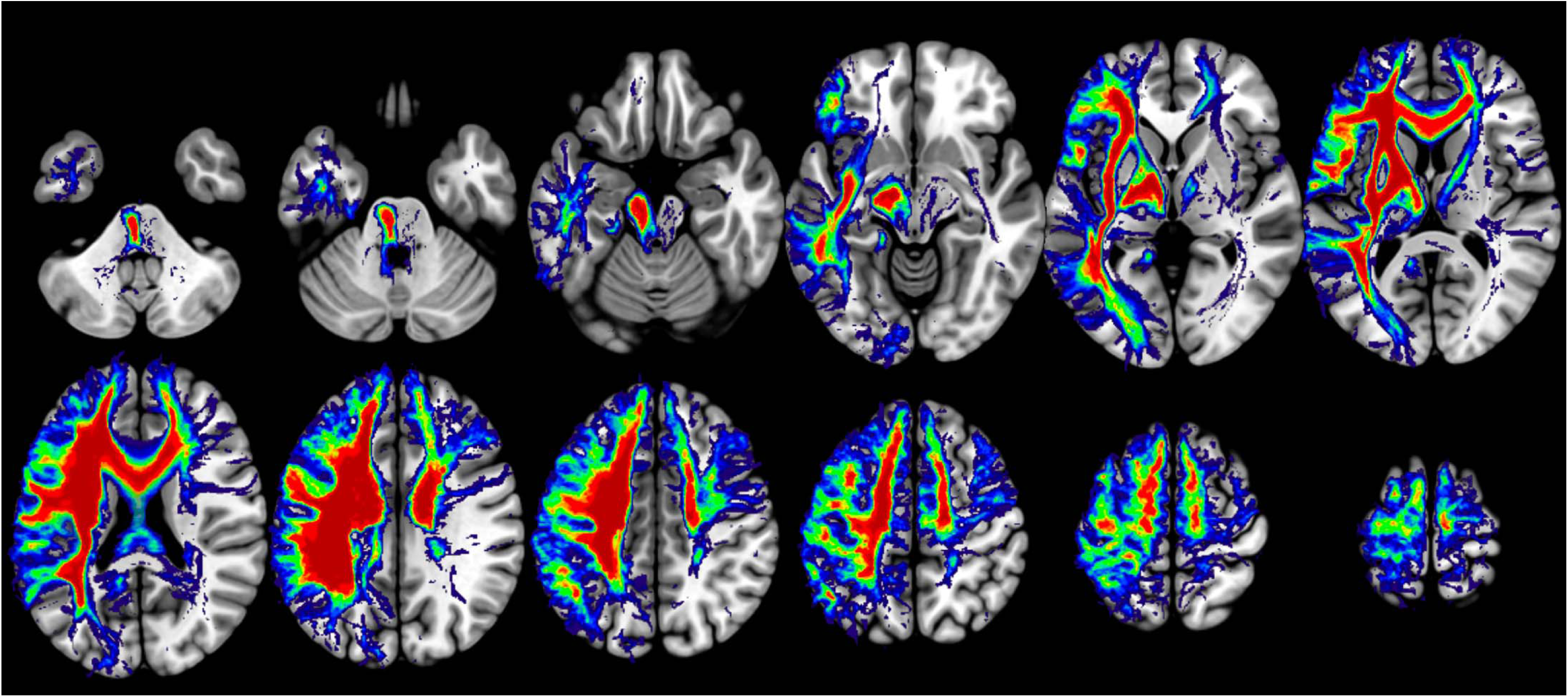
Participant 02 disconnectome map.

**Figure S3.**
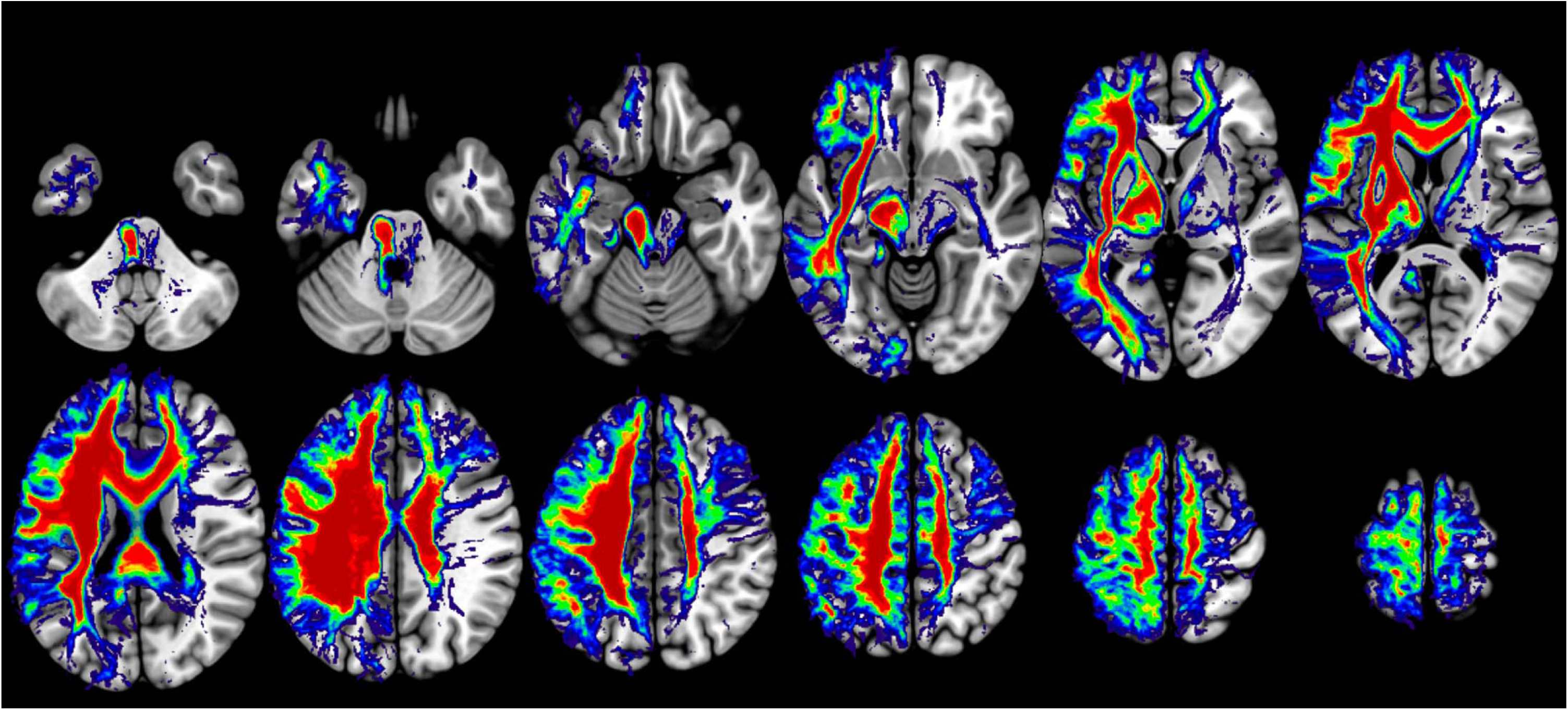
Participant 03 disconnectome map.

**Figure S4.**
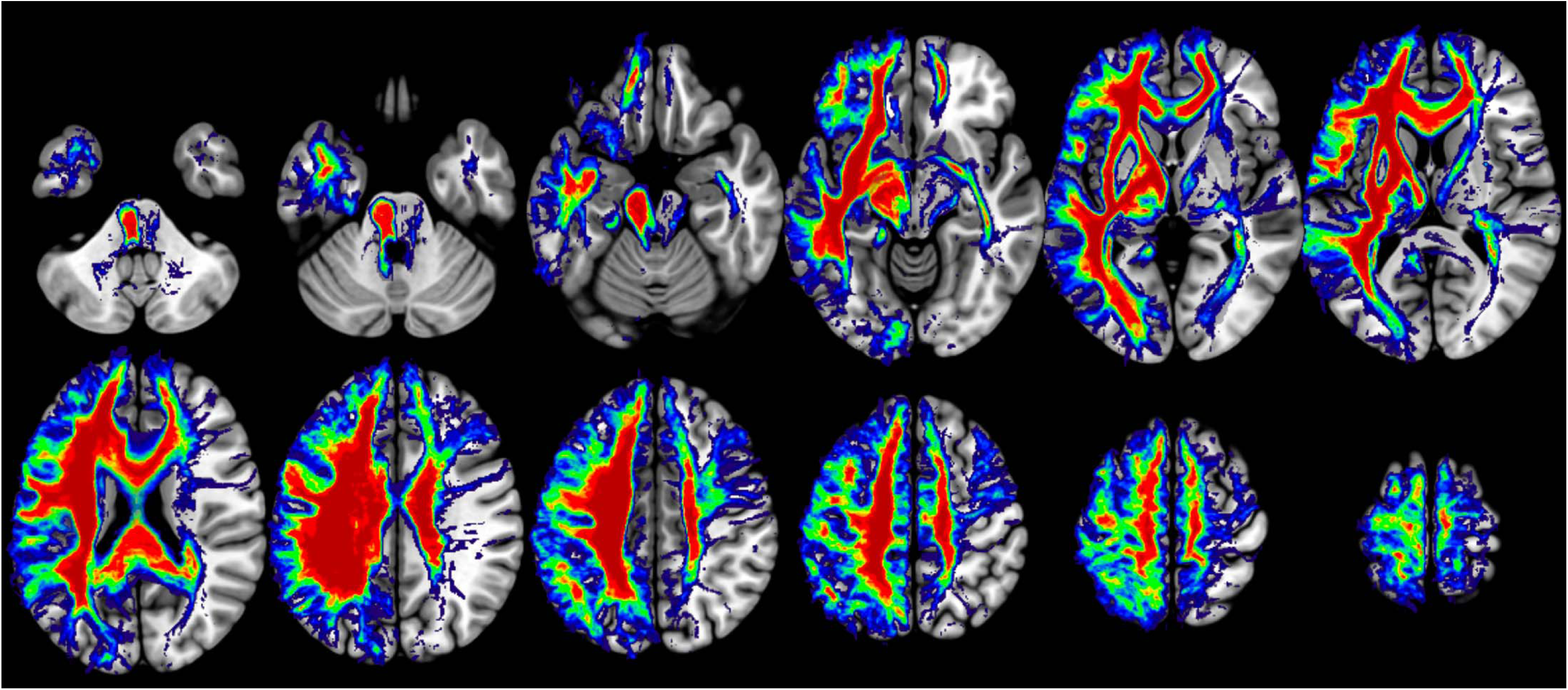
Participant 04 disconnectome map.

**Figure S5.**
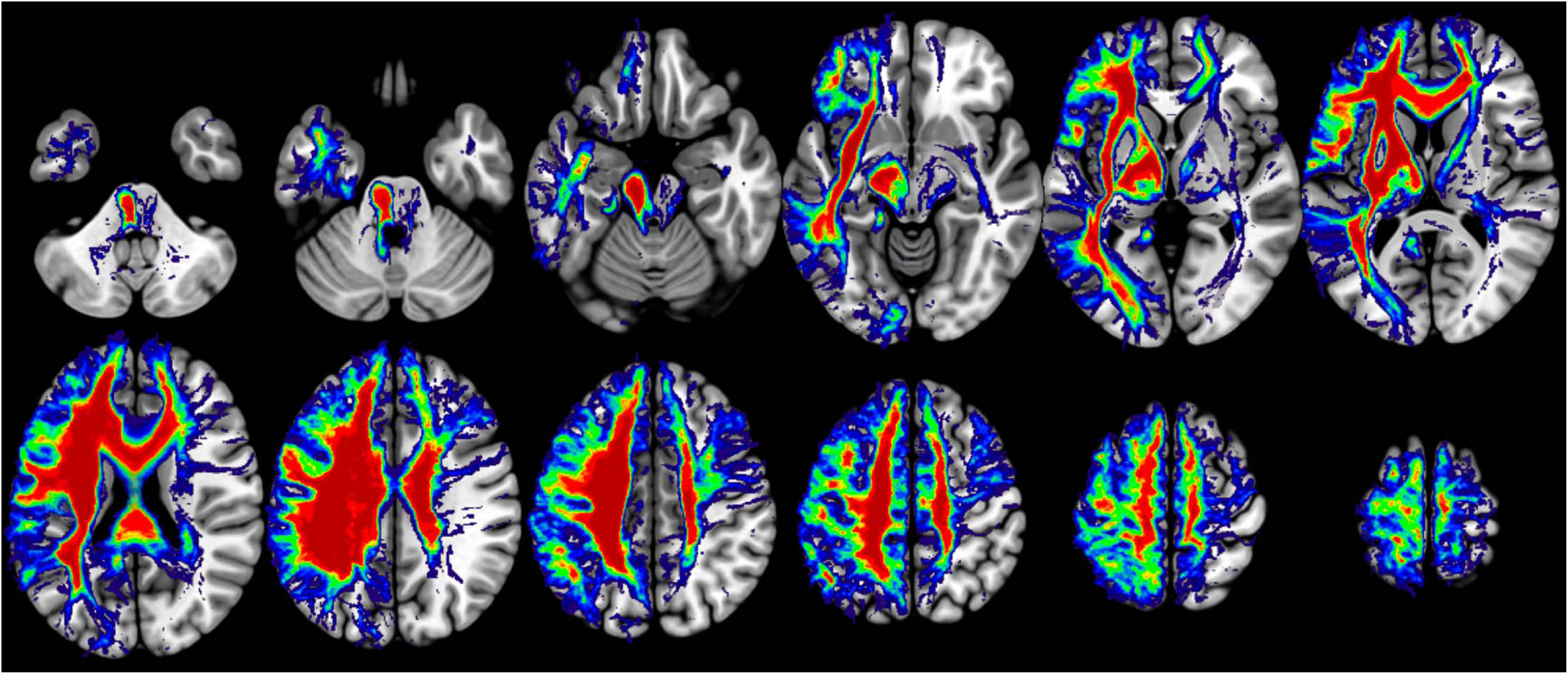
Participant 05 disconnectome map.

**Figure S6.**
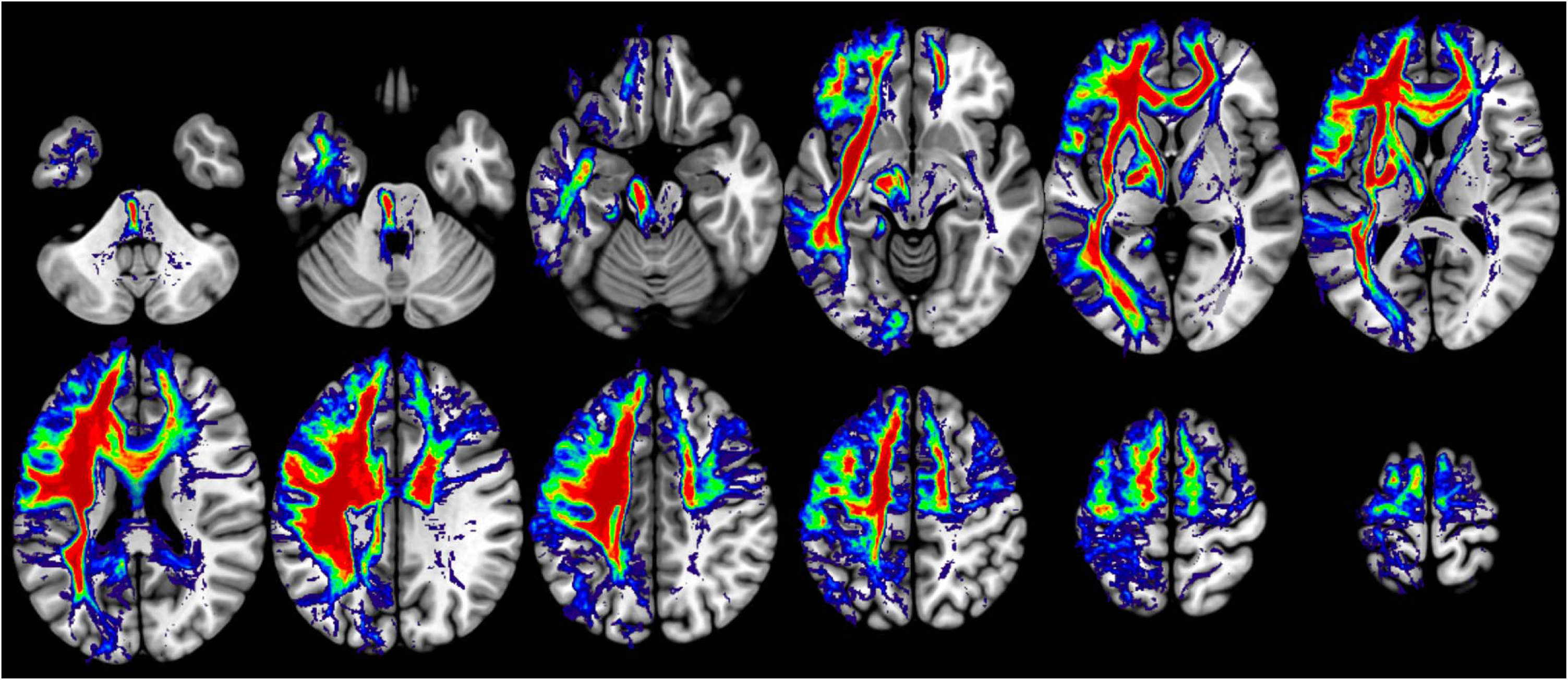
Participant 08 disconnectome map.

**Figure S7.**
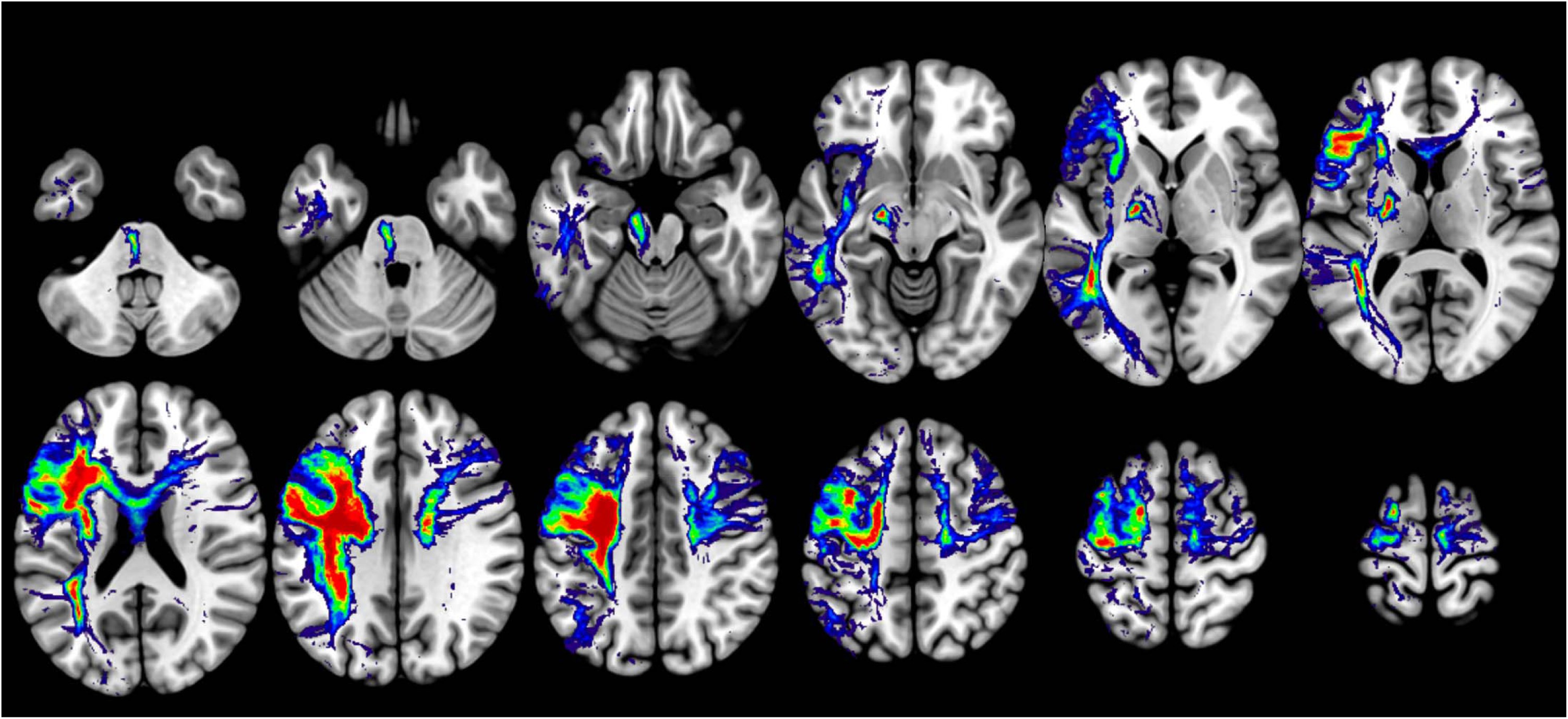
Participant 09 disconnectome map.

**Figure S8.**
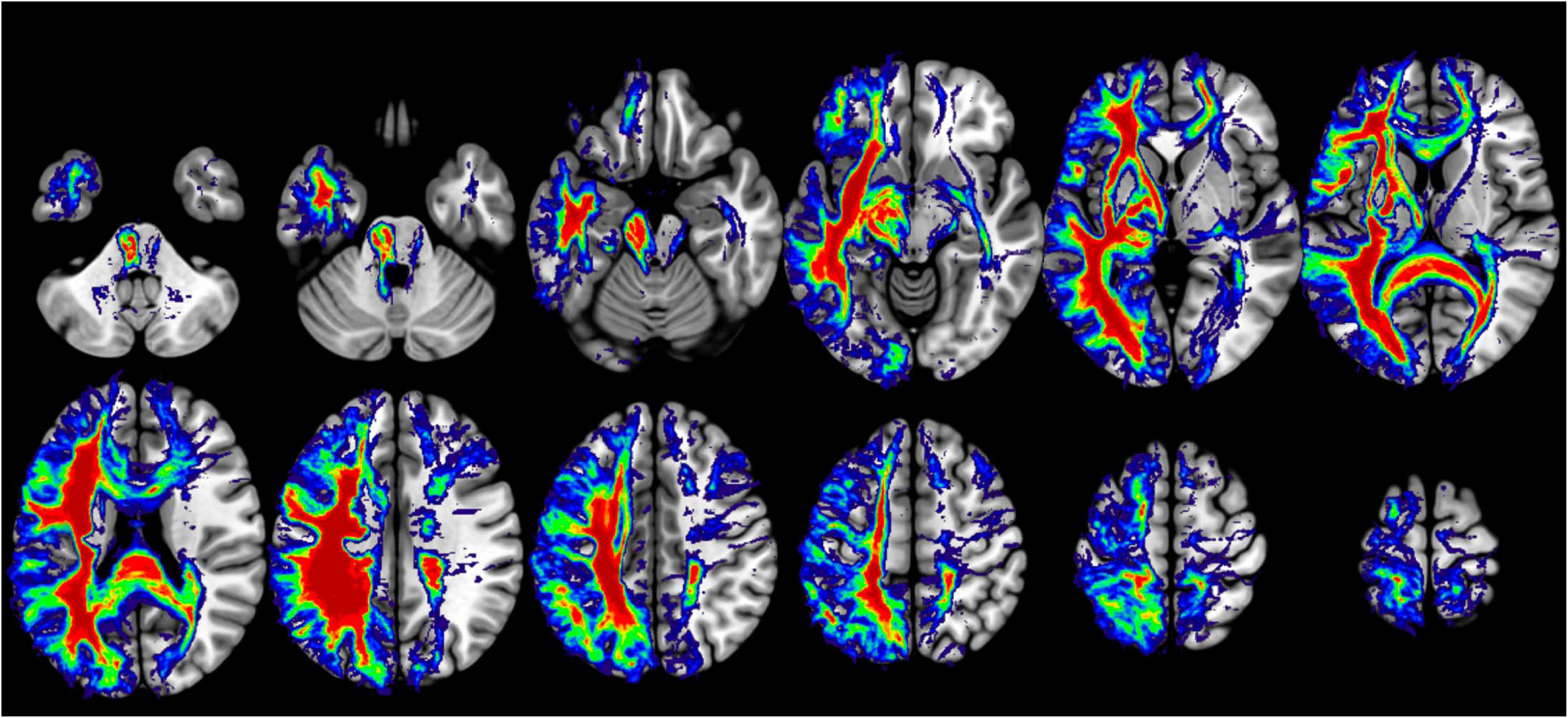
Participant 10 disconnectome map.

